# 3^rd^-ChimeraMiner: A pipeline for integrated analysis of whole genome amplification generated chimeric sequences using long-read sequencing

**DOI:** 10.1101/2022.08.13.503872

**Authors:** Na Lu, Yi Qiao, Pengfei An, Jiajian Luo, Changwei Bi, Musheng Li, Zuhong Lu, Jing Tu

## Abstract

Multiple displacement amplification (MDA) has become one of the most commonly used method of whole genome amplification (WGA) due to the high processivity, strand displacement capacity and high fidelity of the phi29 DNA polymerase, MDA generate vast amount of DNA with higher molecules weight (up to 100kb) and greater genome coverage. Along with the development of the sequencing platform, it is possible to sequence the MDA-amplified DNA molecules with over 20kb by long-read sequencing. However, one of the challenges is the formation of chimeras, which exist in all MDA products, and seriously interfere with the downstream analysis of the long-read sequencing data of MDA-amplified DNA. In this study, we constructed 3^rd^-ChimeraMiner, a chimera detection pipeline for analyzing the long-read sequencing of MDA products, recognizing chimeras, and integrating chimeras into the downstream analysis. Five sequencing data of MDA with different magnification fold were analyzed in here, the proportions of chimeras are much higher than that of next-generation sequencing reads and increase with the increase of magnification folds, ranging from 42% to over 76%. After comparing, 99.92% of recognized chimeras have been demonstrated not to exist in original genomes. After detecting chimeras by 3^rd^-ChimeraMiner, the full-length mapping ratio increased, means more PacBio data could be used in downstream analysis, and mean 97.77% inversions were removed after transferred chimeras into normal reads. 3^rd^-ChimeraMiner revealed efficiency and accuracy in discovering chimeras from long-read sequencing data of MDA, and is promising to be widely used in single-cell sequencing.

## 1. Introduction

The progress of genomic technology has changed the research of molecular biology and diseases treatment strategies^1–6^. In comparing with traditional sanger sequencing technology, short-read next-generation sequencing (NGS) technology^7–9^ offers a great advantage of high throughput^10, 11^, low cost, fast^12^, and high sensitivity^13^, but it exists GC-content bias^14^ and typically only sequences 50-300 base pairs (bp) in length^15^. The advent of the third-generation sequencing (TGS) based on the single molecular technique^16, 17^ overcomes these obstacles based on its capability to read through high GC-content regions and long read length^15^.

Both NGS and TGS need sufficient input DNA to construct sequencing libraries^15^, usually 1-100ng for NGS and 1 μg for TGS^13^. In sometimes, the input DNA is limited, such as single cell sequencing, where whole-genome amplification (WGA) in prior is required^18, 19^. In recent years, several whole genome amplification methods have been developed to operate single cells, such as degenerate oligonucleotide primed PCR (DOP-PCR)^20^, multiple displacement amplification (MDA)^21–24^, and multiple annealing and looping-based amplification cycles (MALBAC)^25^. Among these methods, MDA using phi29 DNA polymerase (phi29DNApol)^26, 27^ and random primers^28^ is the most commonly used method nowadays which has gotten gratifying results in a variety of applications. MDA^28^ generates sufficient quantities of replicated DNA with high fidelity, less amplification bias and large fragment size. Researchers have also developed many improved MDA methods to further optimize the performance, including microwell MDA^29^, emulsion MDA (eMDA)^30^, and microchannel MDA (μcMDA)^31, 32^. However, chimeric sequences (*i.e*. chimeras) ^33^ have been reported in almost sequencing studies using MDA in template preparation^33–35^. Chimeras (Figure 1) are considered as the results of mis-priming during the MDA^33, 36^, appear as DNA rearrangements, cannot be used for genome assembly^34, 37–40^, and make confusion in genetic variation analysis^41–44^. The present of chimeras is an unevadable problem of MDA and has caught more and more attentions^45 46–48^.

**Figure 1.**
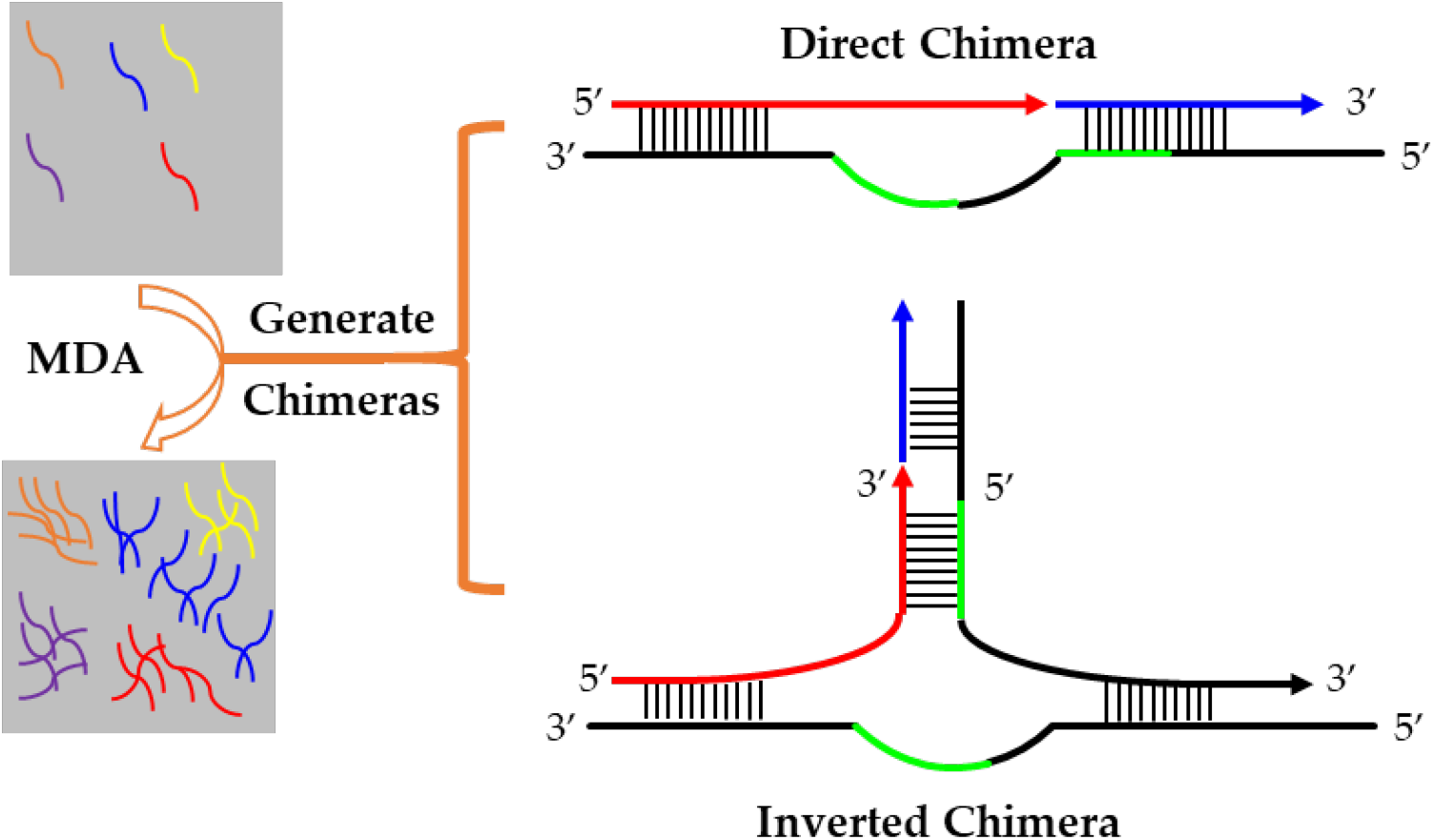
The mechanisms of chimera formation with direct sequences and inverted sequences during the MDA processing. The red lines and the blue lines are the two adjacent segments of a chimera, a mis-priming event occurs during the MDA procedure. The thick black lines are the DNA template strand. The green lines are same sequences on adjacent templates. The thin black lines represent base pairing. There are two types chimera, inverted chimera and direct chimera.

Up till now, most single cell whole genome sequencing (scWGS) studies are based on NGS platforms which generate highly accurate but relatively short reads. There are only a few works perform scWGS using long-read sequencing. But TGS techniques hold many promises in scWGS in future due to their advantages in analyzing genetic variation^49, 50^ and detecting ‘dark regions’ in genome^51^. In the existing works, the proportion of chimeric reads in all MDA NGS reads ranges from 1% to 15%^33, 36, 52–54^. Considering the significantly longer read length, the proportion of chimeric reads in TGS reads is expected to be notably higher, which will make the data analysis more confused. Therefore, a useful pipeline and integrated analysis are desired for whole genome amplification generated chimeric sequences in TGS reads.

In this study, coupled with MDA and PacBio single molecule sequencing platform, we investigate the generation and characteristic of chimeric reads in TGS reads, and established a new bioinformatics pipeline to recognize and classify chimeras in PacBio long reads, called as 3^rd^-ChimeraMiner. In comparing with the ChimeraMiner designed to analyze chimeras in NGS data, 3^rd^-ChimeraMiner is benefit in processing long reads, and 99.92% of recognized chimeric sequences have been demonstrated not to exist in original genomes. By analyzing the TGS reads, the proportions of chimeric reads are much higher than that of NGS reads and increase with the increase of amplification folds, ranging from 42% to over 76%. The results revealed that the mis-priming events in amplification are more frequently occurred than previously known and gradually accumulate with the amplification.

## 2. Results

### 2.1. Design of the 3^rd^-ChimeraMiner

To detect the chimeras in TGS data, a pipeline called 3^rd^-ChimeraMiner was designed and established, which is able to identify valid chimeras from raw PacBio data. The process is shown in Figure 2, in summary, the raw PacBio data was performed quality control by SMRTLink v5.0, the clean PacBio data was aligned to hg19 by using minimap2^55^, the aligned reads was classified as SMRs and CMRs, the SMRs were regarded as candidate chimeras, the segments of the SMRs were obtained and sort based on the alignments, finally the adjacent segments were analyzed and determined by searching the overlap sequence, the valid chimeras were achieved by 3^rd^-ChimeraMiner.

**Figure 2.**
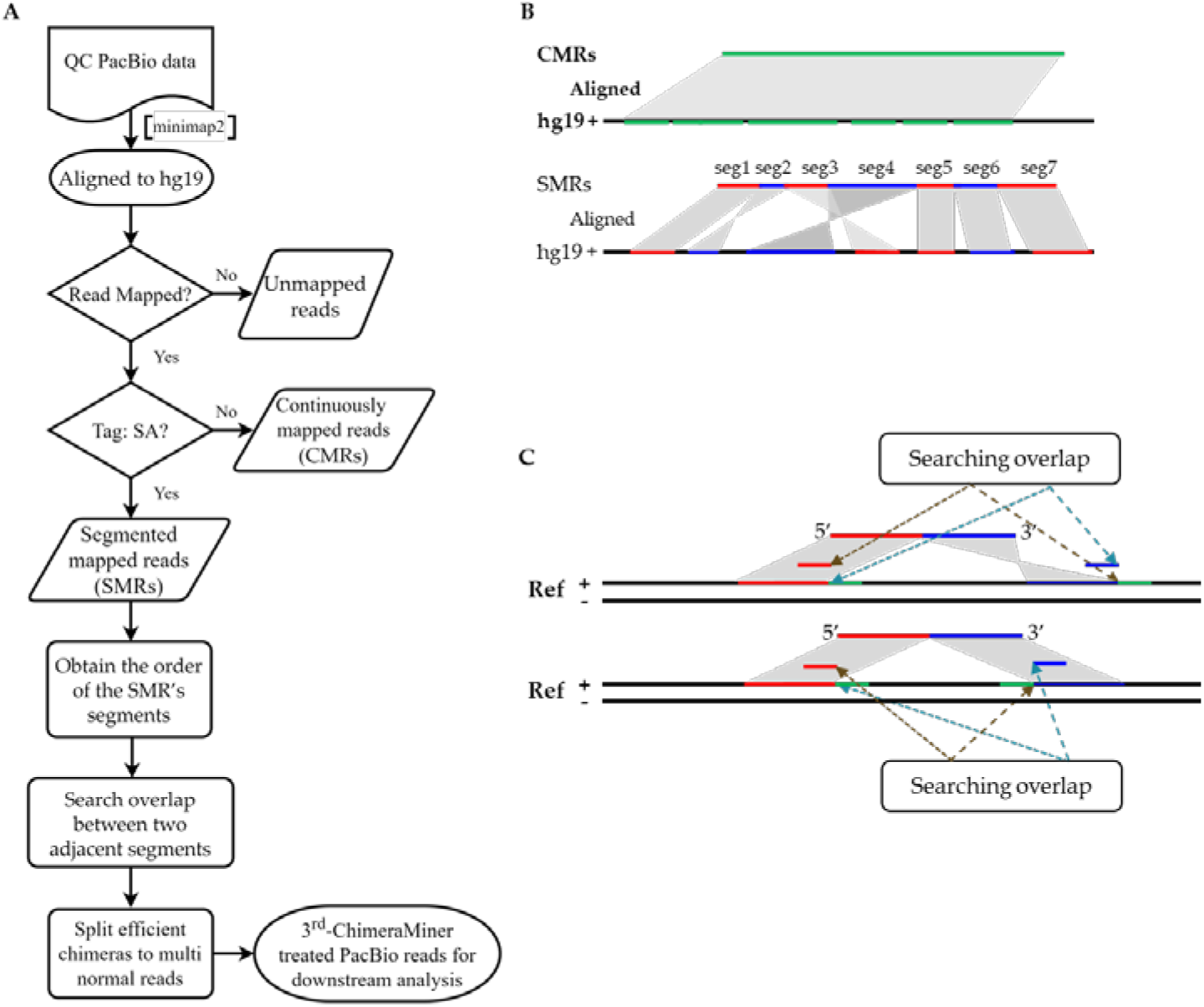
The outline of 3^rd^-ChimeraMiner. A. The bioinformatic analysis pipeline for detecting chimera in TGS data, the detail methods described in Materials and Methods section. B. The detail alignment of CMRs aligned to the hg19 and the detail alignment of SMRs aligned to the hg19. C. Searching overlap between adjacent segments and achieving effective chimeras.

### 2.2. Amplification of micro-quantity DNA with MDA

Five different amounts of genomic DNA obtained from GM12878 cells were used as input for MDA reaction, and the lowest is close to the amount of genomic DNA in a single cell. The MDA reactions generated roughly identical amplified DNA yield (from 19.4 ug to 23.75 ug) and the fragment size distributions displayed peaks around 7.3 ~ 9.2 kb (Table 1). Five MDA samples were selected for PacBio long-read sequencing.

**Table 1:** MDA amplification conditions and yields.

| Sample <sup>1</sup> | Concentration (pg/μL) | Time (h) | Yield (ng/μL) | $\log_{10}(\text{AF})$ <sup>2</sup> | Peak Fragment (bp) |
| --- | --- | --- | --- | --- | --- |
| AF106 | 0.5 | 4 | 388 | 5.89 | 8,041 |
| AF105 | 5 | 4 | 475 | 4.98 | 7,304 |
| AF104 | 50 | 3 | 408 | 3.91 | 8,499 |
| AF103 | 500 | 2 | 432 | 2.94 | 9,084 |
| AF102 | 5,000 | 2 | 436 | 1.94 | 9,186 |

<sup>1</sup> **Sample**, the rules for naming samples, amplification fold referred as AF, and used exponentiation for showing AF, the base is 10 and the exponent or power is n (2, 3, 4, 5, 6). Therefore, AF106 represents the initial DNA of this sample was amplified around 10<sup>6</sup>-fold by using MDA to meet the requirements for PacBio Sequel I library construction. The volume of the whole reaction system of each sample is 50 μL.
<sup>2</sup> **$\log_{10}(\text{AF})$** , AF represents amplification fold, calculated from dividing the DNA Yield by the initial DNA quantity, the AF value is too big that using $\log_{10}$ for showing.

### 2.3 Long-read whole-genome sequencing of micro-quantity DNA and mapping of reads

The libraries of five amplified-MDA sample were constructed and sequenced on the PacBio Sequel I platforms, according to the manufacturer’s instruction. The software SMRTlink v5.0 (--minLength=50, -- minReadScore=0.8) was used for PacBio data cleaning and quality control, about 13 - 19 giga-bases (GB) data were obtained for each five MDA samples (Table 2), and about 3 – 4.5 million subreads after removing the adapter sequences. When we finished the mapping of the subreads and summarized that almost all subreads could be aligned to the hg19 reference, but we found that the full-length mapped ratio relative to mapping ratio is much lower (Table 2). The alignment of each sample was more than the number of subreads, it showed that chimeric reads from the MDA are found in most of PacBio subreads, since each PacBio subreads gives rise to 2–3 separate alignments on average. The aligned read length is a good indicator of the non-chimeric portion of the subreads, as it corresponds to the longest subsequence that can be continuously mapped to the hg19 reference. The N50 aligned read length was ranged from 2 kb for AF106 A to 4 kb for AF102, and the maximum read alignment was ranged from 26.8 kb (AF106) to 62.5 kb (AF102). However, there is a higher level of allelic drop out for PacBio subreads in contrast with the illumina NGS data. For AF102, 29.32% of the genome bases was covered, while AF106 had an even lower genome coverage of 7.84%, the base number of each sample was 13.11 GB. With the decreasing of the initial DNA for amplification, the full-length mapped ratio and the genome coverage decreased sharply (Table 2).

**Table 2:** PacBio Sequel I run statistics and alignment results for five samples.

| Sample | AF106 | AF105 | AF104 | AF103 | AF102 |
| --- | --- | --- | --- | --- | --- |
| QC Subread number | 3,550,863 | 4,322,669 | 4,531,567 | 2,980,163 | 4,075,601 |
| Mean length | 3821 | 3873 | 4082 | 4398 | 4547 |
| Mapping ratio | 98.51% | 98.59% | 98.81% | 98.97% | 98.97% |
| Full-length mapped ratio <sup>1</sup> | 14.47% | 16.35% | 27.27% | 37.08% | 49.41% |
| Number of alignments | 11,443,591 | 13,329,361 | 11,452,239 | 6,500,456 | 7,457,299 |
| Aligned read length N50 | 1,978 | 2,112 | 2,757 | 3,371 | 3,986 |
| Depth <sup>2</sup> | 3.80 | 4.74 | 5.38 | 3.87 | 5.54 |
| Coverage <sup>3</sup> | 7.84% | 14.11% | 21.35% | 25.10% | 29.32% |

<sup>1</sup> **Full-length mapped ratio**, if 90% of the subread can be aligned to the reference genome continuously, which was considered as the full-length mapped subread, the ratio was calculated by dividing the full-length mapped subreads by the number of all subreads.
<sup>2</sup> **Depth**, the mean sequencing depth, the ratio of the total number of sequenced bases to the size of the reference genome.
<sup>3</sup> **Coverage**, here we present the normalized genome coverage, because of the subreads base of AF103 is minimal (13.11 GB), we down-sampled the subreads base other four samples (AF102, AF104, AF105, AF106) close to it, then we calculated the over 1× genome coverage (the proportion of sequences obtained by sequencing to the whole genome) of each sample.

### 2.4 The analysis results from 3^rd^-ChimeraMiner on five MDA samples

Because of the number of alignments was 2–3 fold of the number of subreads, we believed that many of subreads are chimeric reads, so we constructed the 3^rd^-ChimeraMiner for discovering and classifying the chimeras on the five PacBio Sequel I sequencing data (AF106, AF105, AF104, AF103 and AF102). The type and number of chimeras were analyzed and compared. When determining the chimeras, we limited the subsection length no less than 50 bp. Moreover, we set the physical genome distance of two subsections ranging from 25 to 10000 bp, and the overlap sequence length longer than two bases. As for the chimeric strand specificity, this phenomenon not only happened between two reverse strands, but also was observed on the same strand. Following the nomenclature of the previous research, fundamental chimerism could be divided into two types: inverted chimeras and direct chimeras ^33, 36, 54^. Two subsections of inverted chimeras are aligned to two reverse strands, while two subsections of direct chimera could be aligned concordantly to one strand. In additional, although there are insertion chimeras in the NGS reads^36, 54^, but the single-stranded characteristics of the PacBio subreads, so there is no insertion chimeras’ detection in 3^rd^-ChimeraMiner.

We developed 3^rd^-ChimeraMiner to recognize PacBio chimeric reads from MDA samples. We first obtained SMRs by alignment of PacBio reads with each other using minimap2 ^55^ in each sample to detect chimeric reads. Then, we found the overlap segment through the aligning situation of the adjacent subsections. As shown in Table 3, We can find that there is a certain percentage of subreads are chimera in every MDA sample. And for different DNA concertation, the proportion of chimera is not consistent. For PacBio subread, the average chimeric ratio to be 76.16% for AF106 and 42.41% for AF102, indicating a considerably high chimeric ratio in PacBio subreads as observed previously^33–36, 53, 54^. With the increasing of the amplification fold, the chimeric rate and chimeric density increasing simultaneously (Figure 3).

**Table 3:**
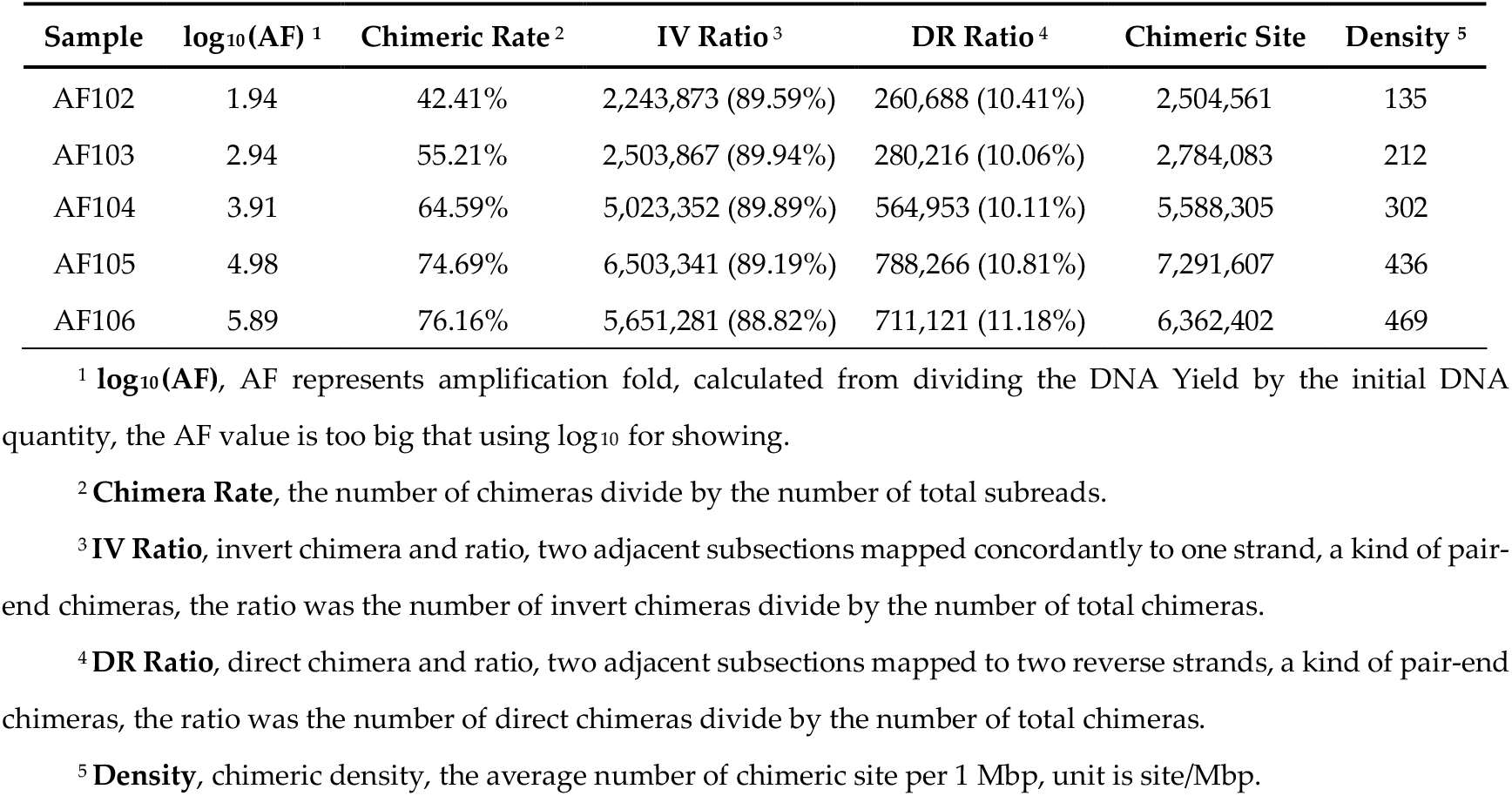
Chimeras analyzing in five MDA samples.

**Figure 3.**
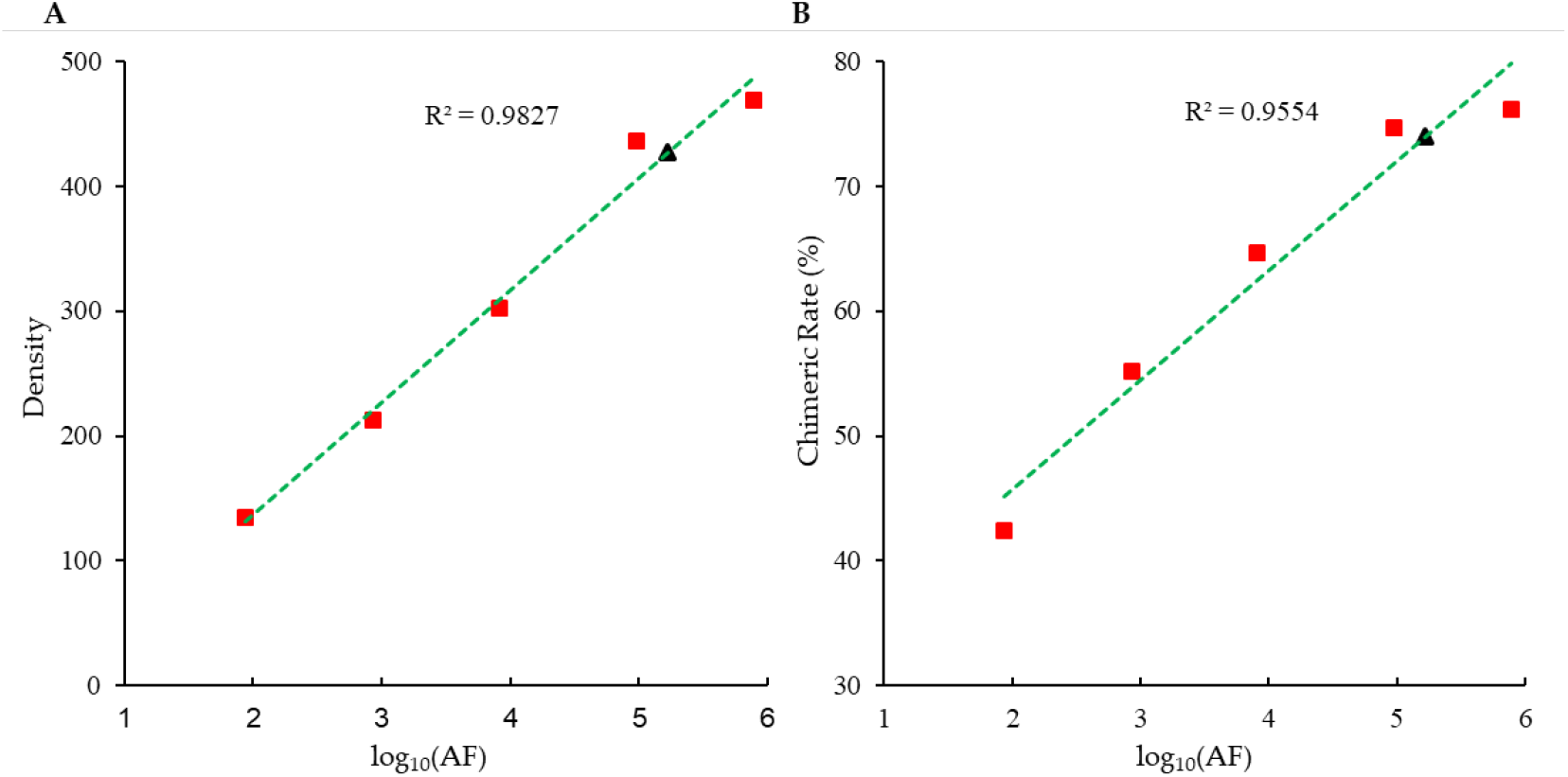
A. The correlation between amplification-fold and chimeric density, the x-axis is the logarithm of the amplification fold, and the y-axis is the number of chimeras on the 1Mbp region. B. The correlation between amplification-fold and chimeric rate, the x-axis is the logarithm of the amplification fold, and the y-axis is the chimeric rate, the number of chimeras divide by the number of total subreads. The red squares are the observed result of five MDA samples (AF102, AF103, AF104, AF105, AF106). The green dotted lines are the second-order polynomial trendline, is obtained by smoothing. We also calculated the amplification fold of sing cell, from 6pg DNA (single cell) to 1ug DNA (the amount of input DNA for PacBio sequencing), the black triangle is the simulation result, it is close to the sample AF105.

Since the segments of the chimeras could locate on either the same strands or the reverse strands, we also finished the quantity statistics of the pair-end chimeras, whose chimeric characteristics were visible, to see the related proportion of these two types. The results illustrated that the chimeric action could be easier to happen between two reverse strands (in about 89.49% for each sample) instead of the same strand (in about 10.51% for each samples), which was generally in concordance with the results of previous researches^33, 35, 36, 54^. We also calculated the occurrence frequency of chimeras in samples with different DNA concentrations by calculating the length of a chimeric event occurred, and the average length of chimera was estimated from 3386bp to 10865bp.

### 2.5 Distribution of the chimeric distance and the overlap length in detected chimeras

In theory, according to the enzyme dynamics of the phi29DNApol, as long as there exists an overlapping section between two chimeric segments, meanwhile the chimeric distance (the genome distance between two chimeric segments) was not very long, there is a probability to generate the chimerism. In order to assess the characteristics of the chimeras, we analyzed the chimeric distance and the length of overlapping section, and derived out the abundance distribution of the five samples (Figure 4.). The results revealed that the abundance of the chimeric distance revealed an approximate bimodal distribution ranging from 0 to 10,000 nt, whose peak values was approximately 750 to 1200 nt, *i.e*. the most probable chimeric distance. Furthermore, the abundance of the overlap length was observed as an approximate decreasing distribution from 3 nt to 30 nt, in which most chimeras had overlap segments was 3 nt.

**Figure 4.**
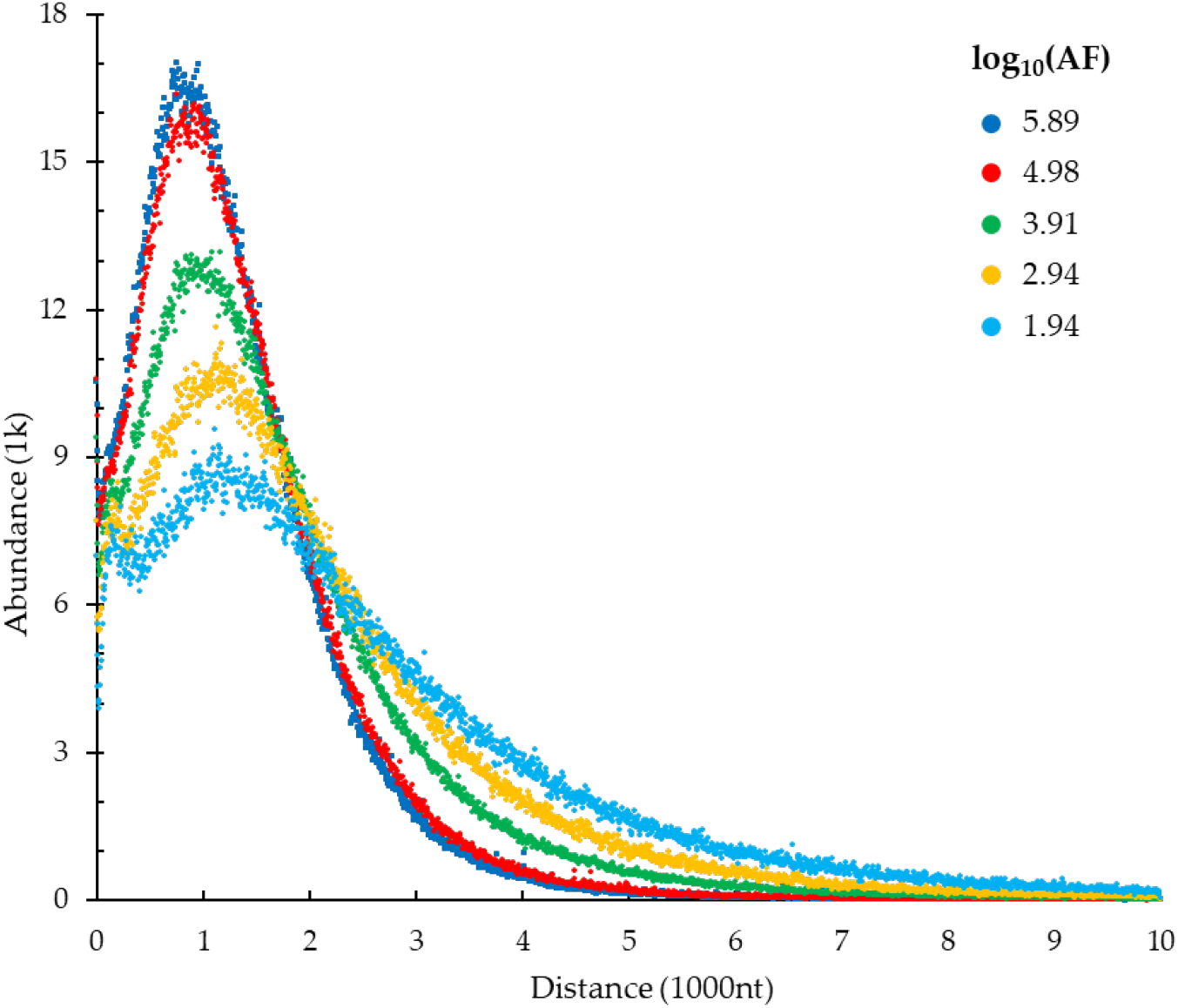
The abundance distribution graph of the physical genomic distance of the two parts of the reads log10(AF), AF is amplification fold, calculated by dividing the DNA Yield by the initial DNA quantity.

### 2.6 The length statistical of subreads, chimeras and non-chimeric subreads on five MDA samples

According to the formation principle of the chimeric sequence, we believed that as the occurrence of chimeric action, a larger DNA fragment will be generated, so that the chimeric subreads and the non-chimeric subreads should have differences in the length of the sequence. In order to compare the length of chimeric subreads and non-chimeric subreads, we calculated the length distribution of each subread type. The results showed in Figure 5 and Table S1. In this figure, we found that the mean length of chimeric subreads is longer than the mean length of non-chimeric subreads. In addition, when the amplification fold increased, we found that the difference of the mean length will be reduced. In here, we presented the results of AF106, the results of several other samples were put inside the appendix. (Figure S2)

**Figure 5.**
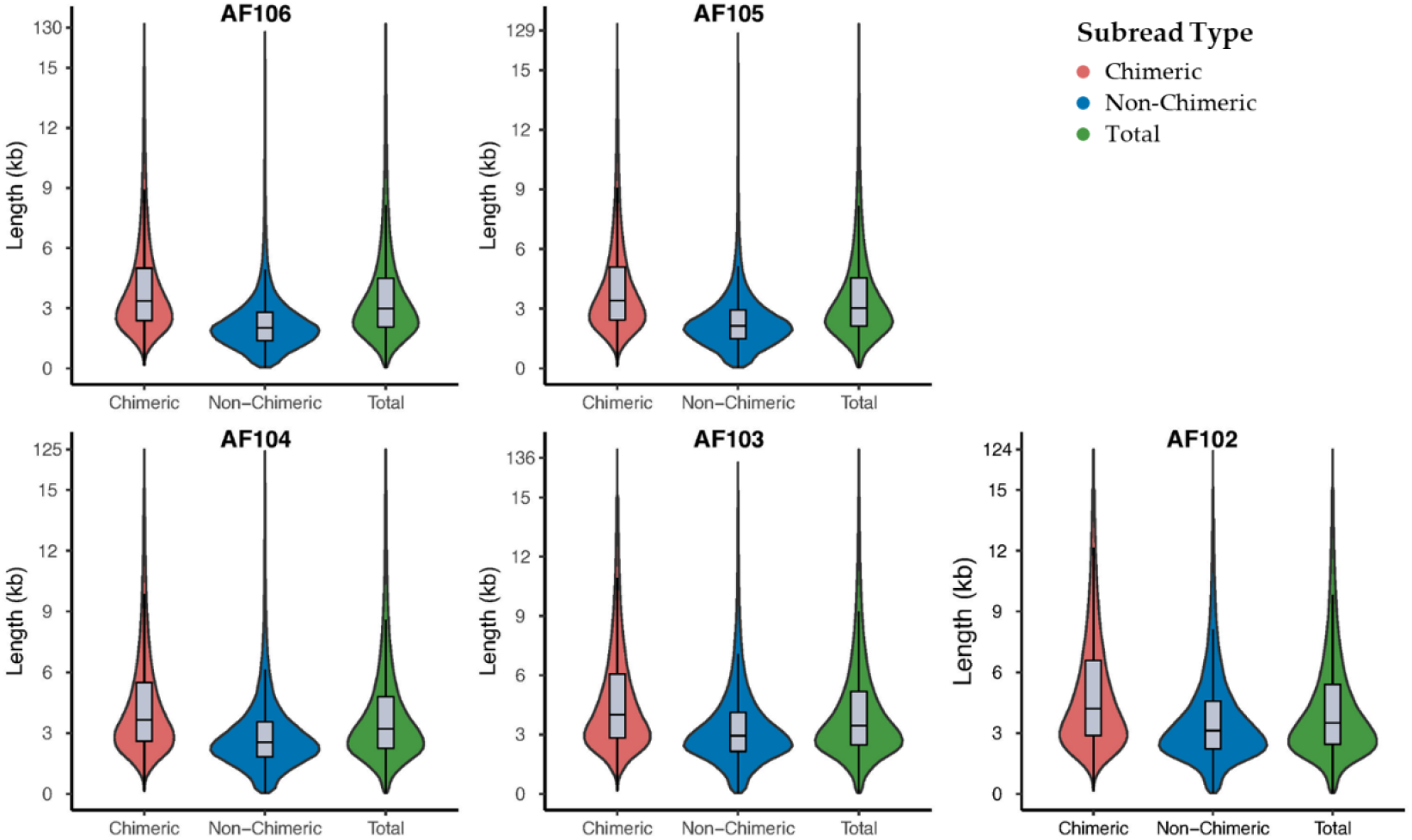
The length distribution of the three subread types (chimeric subread, non-chimeric subread and total subread).

### 2.7 Verifing the accuracy of detected chimeras

In here, we selected the sample AF106, we achieved a reference (length: 1,667,819,669 bp, 6,430,497 contigs, median length is 259 bp) from chimeric reads. The down-sampled dataset contains 619,138,800 150bp reads. After mapped and filtered, we made statistics to the results. We found 0.06% reads (357,821 reads) were mapped to the sequence near the chimeric point, and only 0.08% of the sequence near the chimeric point were covered. The result indicated that 99.92% of the sequence near the chimeric point were impossible to appear in a real sample.

### 2.8 The comparison between chimeras and structural variation

After identifying the chimeric sequences, we extract these subreads from the aligned file, then we split the chimeric subreads into multiple short subreads according to the chimeric point, and then re-aligned them to the reference genome. Merged with the other remained subreads, a new re-aligned bam generated. The five raw aligned bams and five merged aligned bams were used in the downstream structural variations (SVs) analyzing. We also calculated the full-length mapping ratio between raw aligned bams and merged aligned bams in table S2. First, we found that the number of inversions in amplified samples were far greater than that in Bulk sample (mean 400-fold, only 42 inversions in GM12878 Bulk data), there generated many false positive inversions during MDA. After transferred the chimeric reads to normal reads, realigned and SV detection, mean 97.77% of inversions were removed. The results showed in Table S3.

## 3. Discussion

At present, available long-read sequencing platforms provide a powerful approach for clarifying various genomes and promoting the development of genomics. Emerging as a new field of single-cell whole genome sequencing, long-read sequencing platforms have several unique advantages and applications, for example, the precise reference genome assembling of a single cell, the detection and characterization of structural variation are simpler and more effective on the level of single-cell, et al. However, long-read sequencing platforms have limitations on the amount of input DNA (usually requires 1 microgram amount of DNA, equivalent to the amount of genome DNA of hundreds of thousands of human cells) and the use of amplified DNA, especially in the whole genome sequencing of trace samples and single cell samples. Sequencing of intact DNA (as low as 6.6 × 10^−6^ micrograms) from single cell produces a low amount of sequence data which is not enough for analysis. Hence, whole genome amplification (WGS) must be introduced to increase the amount of DNA used for sequencing and multiple displacement amplification (MDA) is the suitable WGA method among many methods. But, MDA is highly susceptible to contamination, especially when working with single cell, and that of amplified DNA with MDA will produce a number of chimeric reads, which are derived from artificial sequences connecting discontinuous DNA regions caused by MDA, and hampers downstream bioinformation analysis (for example proper assembly, full-mapped ratio and structural variation detection, et al). Chimera formation is a challenge in single-cell MDA long read sequencing. To the best of our knowledge, there is no systematic discovery and analysis of chimeras in single-cell MDA PacBio sequencing data previously. Herein, we successfully utilized 3^rd^-ChimeraMiner to distinguish chimeras and normal subreads in the single cell PacBio sequencing data with MDA.

In NGS paired-end (PE) data, the chimeras were initially classified in single-end chimera (located in a read) and paired-end chimera (in the insertion sequence of a pair of reads) by using ChimeraMiner ^54^. But TGS data is single-end, there does not exist paired-end chimera, and we only detected single-end chimeras in these five PacBio data. The single-end chimeras were further divided into direct chimeras and inverted chimeras, based on the orientation of the two adjacent subsections of the chimeric subread. Normally, the single-end chimeras were undoubtedly abandoned in almost studies ^36, 54^. The 3^rd^-ChimeraMiner developed in this study is a novel tool for detecting chimeric reads with high accuracy in PacBio reads from phi29DNApol-mediated MDA amplified products of single cell samples. 3^rd^-ChimeraMiner included several steps: (a) definition of CMRs and SMRs, (b) unscrambling of the SMRs with CIGAR and SA TAG information, (c) searching overlap of adjacent segments of SMRs, (d) showing the efficient chimeras (the details presented in Figure 2 and Materials and methods).

In this study, we used phi29DNApol-mediated MDA for amplifying five samples which the initial concentration of DNA template was 5000, 500, 50, 5, 0.5 pg/μL, and sequenced these samples with PacBio Sequel I. We aligned the PacBio reads to the hg19 with minimap2 ^55^ and performed the statistics of alignment results. In the Table 2, it showed that chimeric reads derived from the phi29DNApol-mediated MDA were found in the most of PacBio reads, since each PacBio read gives rise to between 2 and 3 separate alignments on average in each sample, it needs to be detected and spilt. We used 3^rd^-ChimeraMiner to comprehensively analyze the MDA chimeras in the five phi29DNApol-mediated MDA samples. In addition, we defined the number of chimeric points of 1 Mbp region as the occurrence frequency of chimera, also can regarded as chimeric density. And when we used 3^rd^-ChimeraMiner in single-cell level MDA sample (AF106), the mean chimeric density is 469, it closed to one chimeric sequence was found per 2132 bp of MDA generated DNA in AF106, and the chimeric rate was 76.16%. It indicated that each read of AF106 maybe chimeras. Meanwhile, we found that the chimeric density and chimeric rate is gradually increasing with the increasing of the amplification fold.

After detecting chimeras in all samples, we statistically analyzed the two types of chimeras. First, we found that the AF106 sample generate more chimeras. It is mainly because the amount of input DNA used in AF106 was lower than other samples and amplification fold was the largest. The limited amount of input DNA caused the increase of amplification duration and fold, and the number of chimeras increased synchronously. The possible generation mechanism of the two single-end chimeras in NGS data has been illustrated before in previous studies, and the ratio of these two kinds of chimeras has been calculated. Among PacBio sequencing data, inverted chimera was remaining the dominant type of the single-end chimera (Table 3), which is coherent with the results of NGS sequencing data of MDA and also the previous works ^33, 36, 54^, and the ratio of direct and inverted chimeras was very close (1:9 for 3^rd^-ChimeraMiner and 1.5:8.5 for ChimeraMiner ^54^). The results demonstrated that applying 3^rd^-ChimeraMiner in phi29DNApol-mediated MDA PacBio sequencing data was as effective as in MDA NGS sequencing datasets. All the results indicated that the amplified DNA by using MDA, would generate chimeras, and chimeras will present in most of reads. In addition, the abundance of chimeric distance of chimeras detected by 3^rd^-ChimeraMiner was similar to ChimeraMiner ^54^, an approximate bimodal distribution ranging from 0 to 10000 nt. Based on the results above, 3^rd^-ChimeraMiner showed a promoted efficiency and accuracy in handling the PacBio sequencing data of MDA.

After detected chimeras in PacBio MDA data by using 3rd-ChimeraMiner. We want to know the accuracy of the detected chimeras. We focused on analyzing the sample AF106, we aligned an NGS bulk dataset to the sequences near the detected chimeric points. We found only 0.06% reads (357,821 reads in 619,138,800 reads) were perfect mapped, and only 0.08% of the sequence near the chimeric point were covered. The result indicated that 99.92% of the sequence near the chimeric point were impossible to appear in a real sample, so the chimeras detected by 3^rd^-ChimeraMiner is accurate, and 3^rd^-ChimeraMiner is an efficient pipeline for detecting chimeras from MDA PacBio sequencing data.

In previous studies, we known that the chimera introduced many false positive SVs in amplified samples. In order to prevent the case where the chimeras detected by 3^rd^-ChimeraMiner were true positive structural variations, we have done the following verification analysis. First, we detected SVs in PacBio sequencing data of amplified genomic DNA with MDA and non-amplified genomic DNA (GM12878 Bulk sample) by using pbsv. Used 3^rd^-ChimeraMiner detect chimeras in the five PacBio MDA data. Then we transferred the chimeric reads to normal reads by splitting chimera to multi-subsections based on the chimeric point, and realigned it to the hg19 by using minimap2 ^55^ and performed SV detection by using pbsv (https://github.com/PacificBiosciences/pbsv). After detection, we found that the full-length mapping ratio increased and more PacBio data could be used in downstream analysis. Mean 97.77% inversions were removed after transferred chimeric reads.

In this project, we opted for PacBio sequencing since it currently offers higher per-read accuracy. 3^rd^-ChimeraMiner also be effective to detect chimeras in nanopore WGS data ^56^, but higher error rate of nanopore reads ^56^ will need to allow more mismatch number when searching for overlap sequence.

In conclusion, we demonstrate that long-read sequencing can be performed for single-cell level. We constructed 3^rd^-ChimeraMiner for detecting chimeras from PacBio sequencing data of phi29DNApol-mediated MDA and by applying 3^rd^-ChimeraMiner in several PacBio sequencing data, two single-end chimeras were discovered in per sample dataset. 3^rd^-ChimeraMiner revealed efficiency in discovering chimeric sequences in PacBio data, and is promising to be widely used in single-cell long-read sequencing data to reduce the influence of chimeras to single-cell studies, it may promote near-complete genome assemblies and full haplotype reconstructions from individual cells in the future. In fact, the single-end chimeras could be converted into useful data by splitting from the chimeric sites after detection. Therefore, our study improves the utilization efficiency of the PacBio sequencing data, and is able to integrate the single-end chimeras into the following bioinformatics analysis, rapidly.

## 4. Materials and Methods

### 4.1 Micro DNA of whole-genome amplification by regular MDA

Genomic DNA was extracted from GM12878 cell line with a DNA extraction kit (QIAamp DNA Mini Kit, QIAGEN). The extracted DNA was then diluted sequentially and amplified with modified thermotolerant phi29 DNA polymerase (EquiPhi29 polymerase, Thermo Scientific). For the 5 reactions, the initial concentration of DNA was set to 5,000, 500, 50, 5 and 0.5 pg/μL, and the reaction time was 2, 2, 3, 4 and 4h, respectively. Each mixed solution of MDA contained 25 μM 6N primer (Sangon, China), 4 mM dNTP (NEB, USA), extracted DNA, 1 mM DTT (Thermo Scientific, USA), phi29 DNA polymerase with corresponding buffer at recommended final concentration and nuclease-free water up to 50 μL.

The products were quantified using a Qubit dsDNA kit (Thermo Scientific) after diluted by 20 times. Magnification times were calculated accordingly.

### 4.2 PacBio whole genome sequencing

The libraries of five amplified-MDA sample were constructed and sequenced on the PacBio Sequel I platforms, according to the manufacturer’s instruction. The software SMRTlink v5.0 (--minLength=50, --minReadScore=0.8) was used for PacBio data cleaning, quality control and removing the adapter sequences.

### 4.3 Bioinformatics Analysis for chimera detecting in PacBio data

Using the pipeline of previous studies as guideline, 3^rd^-ChimeraMiner was composed of three parts: A. sequencing reads pre-processing, reads alignment and filtrating; B. unscrambling of the soft-clipped alignment reads and realignment; C. Searching overlap and achieving valid chimeras.

#### 4.3.1. Pre-processing, alignment and filtrating

Firstly, we achieved the clean subreads through removed the reads which read length below 50 bp. The clean subreads were then mapped to the hg19 human reference genome by using minimap2^55^. The mapped data was saved as BAM file with SAMtools ^57^ view (1.11-5-g0920974) and then statistically analyzed through Sambamba ^58^ and QualiMap2 ^59^ bamqc (v.2.2.2-dev), and the aligned detail information was explored. By analyzing the SAM “SA” tag and SAM FLAG value, we define those mapped reads which no “SA” tag as continuously mapped reads (CMRs), and define the mapped reads which have “SA” tag as partially mapped reads (SMRs), SMRs will appear in alignment result many times according to the soft-clipped alignments and are considered candidate chimeric reads. Here, the SMRs were retained for the following analysis by unscrambling the “SA” CIGAR information.

#### 4.3.2. Unscrambling of the SMRs

In this step, firstly, we determine the position (start and end) of each segment of the retained SMRs in the original read in accordance with the soft-clipped alignment locations, strand orientation of mapped and mapped order. Then, we split the SMRs to multiple segments according to the positions. The number of split segments was calculated through following equation:

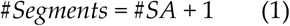

When using Minimap2 for mapping reads to reference genome, if 5 parts of one PacBio read can be aligned to 5 adjacent but discontinuous regions on one chromosome of hg19, the id of this read will appear 5 times, and it means this read have one best alignment and 4 secondary alignment. Here, we will split this read to 5 segments and the detail information of alignment of each segment also saved for searching the overlap of the adjacent segments.

#### 4.3.3. Searching overlap and achieving effective chimeras

In this step, we aimed to search the overlap sequences of the adjacent split segments and achieve the chimeric sites of all the valid chimeras. Firstly, according the results of alignment, adjacent segments whose two segments mapped to different strands of the chromosome were regarded as potential inverted chimeras and that mapped to same strands of the chromosome were regarded as potential direct chimeras. Because of the not high base accuracy (~85%) of PacBio Sequel I, we designed a different algorithm for searching the overlap sequences of the potential chimeras, which relative to the searching algorithm of ChimeraMiner for NGS. The detail of the algorithm is as described below.

First, we followed the location of the two adjacent segments (AS) in the reference genome, and the adjacent segments which located on different chromosome were excluded. Because it is known from previous studies that the overlap length of the chimeras is mainly located at 3 to 30 bases, so we restricted the length of the overlap sequence of one side with the maximum 30 bp. Then, at the coordinate position of the end of the former subsection of the AS, 31-nt sequences were cut forward on the genome and was named as Former 1 (F1), and 31-nt sequences were cut back on the genome and was named as Former 2 (F2). In addition, at the coordinate position of the front of the latter subsection of the AS, 31-nt sequences were cut forward on the genome and was named as Latter 1 (L1), and 31-nt sequences were cut back on the genome and was named as Latter 2 (L2). If the length of the F1 or L1 was shorter than 31-nt, we polished the subsection with N forward. And if the length of the F2 or L2 was shorter than 31-nt, we polished the subsection with N backward. Next, the F1 was cyclically aligned with the L1 by shortening one nucleotide in one loop from back to front, till the overlap sequence was found or no sequence left. The F2 also was cyclically aligned with the L2 by shortening one nucleotide in one loop from front to back. We tolerated 20% mismatch during the cyclically alignment, the mismatch cannot appear at the beginning of the overlap sequence.

The final overlapping sequence was composed of the sequences found by the above pipeline. We retained the potential chimeras with overlapping sequence length greater than 3 bp and considered them as the effective chimeras.

### 4.4 Verifying the accuracy of the chimeras

Although we recognized the chimeras in PacBio subreads according to the characteristics of the chimeric sequence, the chimeric sequence, but chimeras can generate randomly on all DNA fragments, so we don’t know whether the identified chimeras are accurate. So, we designed a method for analyzing the accuracy of 3^rd^-ChimeraMiner through introduce an NGS bulk dataset compared with sequences near the chimeric point. First, we cut the 260 bp sequence around the chimeric point from the chimera. Then transferred the sequences to a fasta file as a reference and indexed by using BWA^60^. In addition, we down-sampled a dataset contains 30X genome data from a NA12878 Bulk sequencing NGS data (150bp paired-end sequenced by laboratory in previous study). Then we mapped the down-sampled reads to the reference that we achieved in first step by using BWA-MEM. In final, we removed unmapped reads and imperfect aligned reads. We regarded the reads: a. over 145 bp mapped, b. the number of indels less than 2 and no soft-clipped, c. or no indels and the length of soft-clipped less than 5 bp, as imperfect aligned reads. We calculated the covered region of the reference and mapping ratio.

### 4.5 Detecting SV in TGS bulk data and in PacBio MDA data

In addition, in many studies, the generation of chimeric sequences will introduce many false-positive SVs to affect the accurate identification of SV. Therefore, in order to verify the accuracy of the identified chimeras, we built the benchmark SVs of NA12878, which identified SVs of bulk sample (PacBio SequelII CCS sequencing data, downloaded from GIAB FTP) by using pbsv (v2.8.0, https://github.com/PacificBiosciences/pbsv) with default parameters. For raw aligned bam and the five merged aligned bams, we identified SVs by using pbsv (v2.8.0, https://github.com/PacificBiosciences/pbsv) with default parameters. For each called SVs results, we removed the SVs which located in the regions of centromere and telomere. For benchmark SVs, the minimum supporting reads was set to 10. Because of the genome coverage of other sequenced samples in this study were relatively low, so we used the mean depth divided by 1X genome coverage as the minimum supporting reads (55, 33, 22, 17, 16 corresponding to AF106, AF105, AF104, AF103, AF102). In addition, in these callsets, we only focused on the SVs on the autosomes and sex chromosomes and summarized the variations ≥ 50 bp, meanwhile SVs with the type “INV” (inversion) was included. Compare bulk SVs with the valid chimeras and the filtrated invalid chimeras, we verify the effective of the 3^rd^-ChimeraMiner in detecting chimeras from PacBio long-reads.

### 4.6 Chimeric reads were transferred to normal reads

According to the chimeric point detected by using 3^rd^-ChimeraMiner, we split the chimeric long reads to segments and rebuild them to new split reads, save with fastq file. Then, we mapped the rebuilt long reads to hg19 by using the minimap2, the mapped data was saved as BAM file with SAMtools view and then statistically analyzed through Sambamba and QualiMap2 bamqc. We calculate the full-length mapping ratio after processing with the split chimeric reads.

## Author Contributions

J.T. and Z.L. conceived and designed the experiments; Y.Q. performed the experiment. N.L., and A.P. performed the bioinformatics analysis; N.L., J.L and C.B. performed the data statistic and figure plot; N.L., and M.L. contributed analysis tools; J.T. and N.L. wrote the manuscript.

## Acknowledgments

We thank the Big Data Center of Southeast University for providing the facility support on the numerical calculations in this paper. The PacBio MDA sequencing data of this study will be deposited in the Sequence Read Archive (SRA) in future.

## Code availability

The bioinformatics pipeline and all custom codes have been deposited in the GitHub repository (https://github.com/dulunar/3rdChimeraMiner).

