## Supplemental Figures and Tables for "3^rd^-ChimeraMiner: A pipeline for integrated analysis of whole genome amplification generated chimeric sequences using long-read sequencing"

### Supplementary Figures

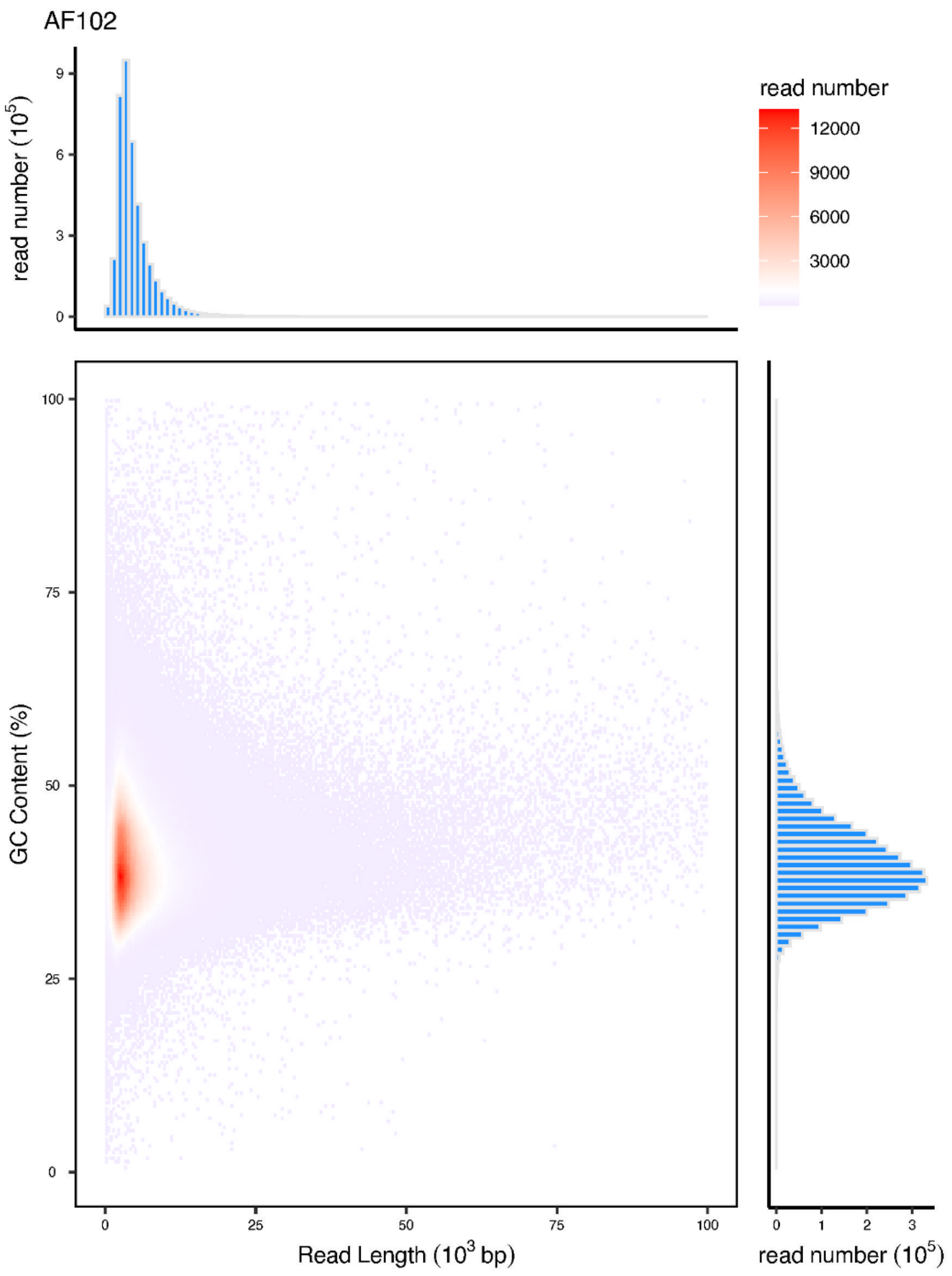

Figure S1. The length and GC content distributions of sequencing reads of AF102 produced from PacBio Sequel I continuous long-read (CLR) technology. Amplification fold is close to 100.

Figure S2. The length and GC content distributions of sequencing reads of AF103 produced from PacBio Sequel I continuous long-read (CLR) technology. Amplification fold is close to 1000.

Figure S3. The length and GC content distributions of sequencing reads of AF104 produced from PacBio Sequel I continuous long-read (CLR) technology. Amplification fold is close to 10000.

Figure S4. The length and GC content distributions of sequencing reads of AF105 produced from PacBio Sequel I continuous long-read (CLR) technology. Amplification fold is close to 10000.

Figure S5. The length and GC content distributions of sequencing reads of AF106 produced from PacBio Sequel I continuous long-read (CLR) technology. Amplification fold is close to 100000.

**Supplementary Tables**

Table S1: Length and number of three subread types

| Sample | Subread Type | Mean Length | Mode Length | Number |
|---|---|---|---|---|
| **AF106** | Total | 3,820 | 2,201 | 3,550,863 |
| | Non-Chimeric | 2,442 | 1,895 | 846,519 |
| | Chimeric | 4,252 | 2,426 | 2,704,344 |
| **AF105** | Total | 3,873 | 2,315 | 4,322,669 |
| | Non-Chimeric | 2,542 | 1,939 | 1,093,941 |
| | Chimeric | 4,324 | 2,471 | 3,228,728 |
| **AF104** | Total | 4,082 | 2,436 | 4,531,567 |
| | Non-Chimeric | 3,042 | 2,184 | 1,604,615 |
| | Chimeric | 4,651 | 2,927 | 2,926,952 |
| **AF103** | Total | 4,398 | 2,631 | 2,980,163 |
| | Non-Chimeric | 3,522 | 2,473 | 1,334,910 |
| | Chimeric | 5,108 | 2,631 | 1,645,253 |
| **AF102** | Total | 4,546 | 2,743 | 4,075,601 |
| | Non-Chimeric | 3,871 | 2,542 | 2,347,215 |
| | Chimeric | 5,463 | 3,041 | 1,728,386 |

Total, all PacBio reads of each sample. Non-Chimeric, the PacBio reads which could be continuously aligned to the reference hg19, or the unmapped PacBio reads. Chimeric, the PacBio reads which only partially be aligned to the reference hg19.

Table S2. Full-length mapping ratio before and after transferred chimeric reads

| Sample | Raw Reads | Transferred Reads |
|---|---|---|
| **AF106** | 14.47% | 79.92% |
| **AF105** | 16.35% | 80.93% |
| **AF104** | 27.27% | 83.69% |
| **AF103** | 37.08% | 85.44% |
| **AF102** | 49.41% | 86.91% |

Full-length mapping reads means the PacBio reads which could be continuously aligned to the reference hg19. Raw Reads, before detection of chimeras in PacBio reads. Transferred Reads, after detection of chimeras in PacBio reads, then the chimeric reads were transferred to normal reads and re-aligned to the reference hg19.

Table S3. SV detection before and after transferred chimeric reads

| INV | AF106 | AF105 | AF104 | AF103 | AF102 |
|---|---|---|---|---|---|
| **Raw reads** | 6769 | 17518 | 25089 | 16438 | 18954 |
| **Transferred reads** | 181 | 622 | 584 | 228 | 232 |
| **Reduction** | 97.33% | 96.45% | 97.67% | 98.61% | 98.78% |

Raw Reads, before detection of chimeras in PacBio reads, aligned to the reference hg19 and detected SV using pbsv. Transferred Reads, after detection of chimeras in PacBio reads, then the chimeric reads were transferred to normal reads and re-aligned to the reference hg19, then merged with the non-chimeric reads and detected SV using pbsv.
